# Cell-autonomous regulation of astrocyte activation by the circadian clock protein BMAL1

**DOI:** 10.1101/362814

**Authors:** Brian V. Lananna, Collin J. Nadarajah, Mariko Izumo, Michelle R. Cedeño, David D. Xiong, Julie Dimitry, Chak Foon Tso, Celia A. McKee, Percy Griffin, Patrick W. Sheehan, Jeffery A. Haspel, Ben A. Barres, Shane A. Liddelow, Joseph S. Takahashi, Ilia N. Karatsoreos, Erik S. Musiek

## Abstract

Circadian clock dysfunction is a common symptom of aging and neurodegenerative diseases, though its impact on brain health is poorly understood. Astrocyte activation occurs in response to diverse insults, and plays a critical role in brain health and disease. We report that the core clock protein BMAL1 regulates astrogliosis in a synergistic manner via a cell-autonomous mechanism, and via a lesser non-cell-autonomous signal from neurons. Astrocyte-specific *Bmal1* deletion induces astrocyte activation *in vitro* and *in vivo*, mediated in part by suppression of glutathione-s-transferase signaling. Functionally, loss of *Bmal1* in astrocytes promotes neuronal death *in vitro*. Our results demonstrate that the core clock protein BMAL1 regulates astrocyte activation and function *in vivo*, elucidating a novel mechanism by which the circadian clock could influence many aspects of brain function and neurologic disease.

**Highlights:**

- Circadian disruption promotes astrocyte activation.
- Astrocyte-specific deletion of the circadian clock gene BMAL1 induces astrocyte activation.
- BMAL1 regulates astrocyte activation by altering glutathione-s-transferase signaling.
- Loss of astrocyte BMAL1 enhances neuronal cell death in a co-culture system.

**eTOC blurb:** Lananna et al. show that the circadian clock protein BMAL1 regulates astrocyte activation via a cell autonomous-mechanism involving diminished glutathione-s-transferase signaling. This finding elucidates a novel function of the core circadian clock in astrocytes, and reveals a BMAL1 as a modulator of astrogliosis.

## Introduction

Changes in behavioral circadian rhythms are common in a wide range of neurodegenerative diseases (Breen et al., 2014; Hatfield et al., 2004; Morton et al., 2005; Musiek and Holtzman, 2016). Studies suggest that such changes can occur early in disease progression (Breen et al., 2014; Hatfield et al., 2004), sometimes before the onset of any other overt symptoms (Musiek et al., 2018; Tranah et al., 2011). Circadian rhythms are generated by the suprachiasmatic nucleus of the hypothalamus (SCN) (Mohawk et al., 2012), which synchronizes cellular clocks throughout the body to the light-dark cycle. The core molecular clock consists of a transcriptional-translational feedback loop with a positive transcriptional limb, consisting of heterodimers of the bHLH transcription factor BMAL1 (*aka* ARNTL, aryl hydrocarbon receptor nuclear translocator-like protein 1) with either CLOCK (circadian locomotor output cycles kaput) or NPAS2 (neuronal PAS domain protein 2). The negative limb includes Period (PER), Cryptochrome (CRY), and REV-ERB genes which are transcriptional targets of BMAL1 and provide negative feedback inhibition of BMAL1 function (Mohawk et al., 2012). The core clock regulates transcription of thousands of genes in a highly tissue-specific manner, regulating cellular functions including metabolism, inflammation, and redox homeostasis (Bass and Takahashi, 2010; Zhang et al., 2014a). In addition to disrupted activity and sleep rhythms, dysregulated circadian gene expression patterns are also observed in neurodegenerative disease models (Song et al., 2015; Stevanovic et al., 2017) and patients (Breen et al., 2014; Cronin et al., 2017). However, the impact of clock dysfunction on neurologic diseases of the brain remains poorly understood.

Astrocytes play a critical role in brain health and neurodegenerative disease (Pekny et al., 2016). Dysfunctional astrocytes can drive neurodegeneration (Lian et al., 2015; Macauley et al., 2011; Yamanaka et al., 2008), while optimization of certain astrocyte functions can protect against neurotoxic stimuli (Kraft et al., 2004; Xiao et al., 2014). Astrocyte activation is a ubiquitous response to brain injury, from neurodegeneration to trauma, which has historically been characterized by increased expression of the cytoskeletal protein glial fibrillary acidic protein (GFAP) (Sofroniew, 2014). Recent work has begun to elucidate the diversity of astrocyte activation phenotypes beyond GFAP, as astrocytes activated by different stimuli express unique transcriptional profiles which can be associated with divergent phenotypes, ranging from neurotoxic to neurotrophic (Liddelow et al., 2017; Zamanian et al., 2012). However, a further understanding of astrocyte activation mechanisms is needed in order to effectively target protective responses in astrocytes therapeutically, while minimizing detrimental processes.

Astrocyte activation and circadian clock dysfunction are two pervasive, often co-existent features of neurological diseases, though their interaction is unknown. We previously observed that deletion of the core clock gene *Bmal1*, which abrogates all circadian clock function, caused severe, spontaneous astrogliosis throughout the mouse brain, which was accompanied by increased oxidative stress, synaptic damage, and inflammation (Musiek et al., 2013). However, the cellular and molecular mechanisms linking BMAL1 to astrocyte activation and function remain uncertain. Thus, we sought to characterize the astrocyte activation induced by circadian clock disruption and evaluate whether the clock protein BMAL1 might regulate astrogliosis in a cell-autonomous manner.

## Results

We first asked if non-genetic disruption of circadian rhythms could influence *Gfap* levels in the brain. Exposure of wild type (WT) C57Bl/6 mice to a 10hr:10hr light:dark cycle induces circadian desynchrony and eventually arrhythmicity, as well as loss of detectable circadian transcript oscillations in the cerebral cortex (Fig. 1A and S1A). After 6 weeks of circadian disruption, *Gfap* transcript levels were increased throughout the circadian cycle (Fig. S1B), with an average increase of 19.7% (Fig. 1B). Thus, light-induced behavioral circadian rhythm disruption promotes astrocyte activation.

**Figure 1.**
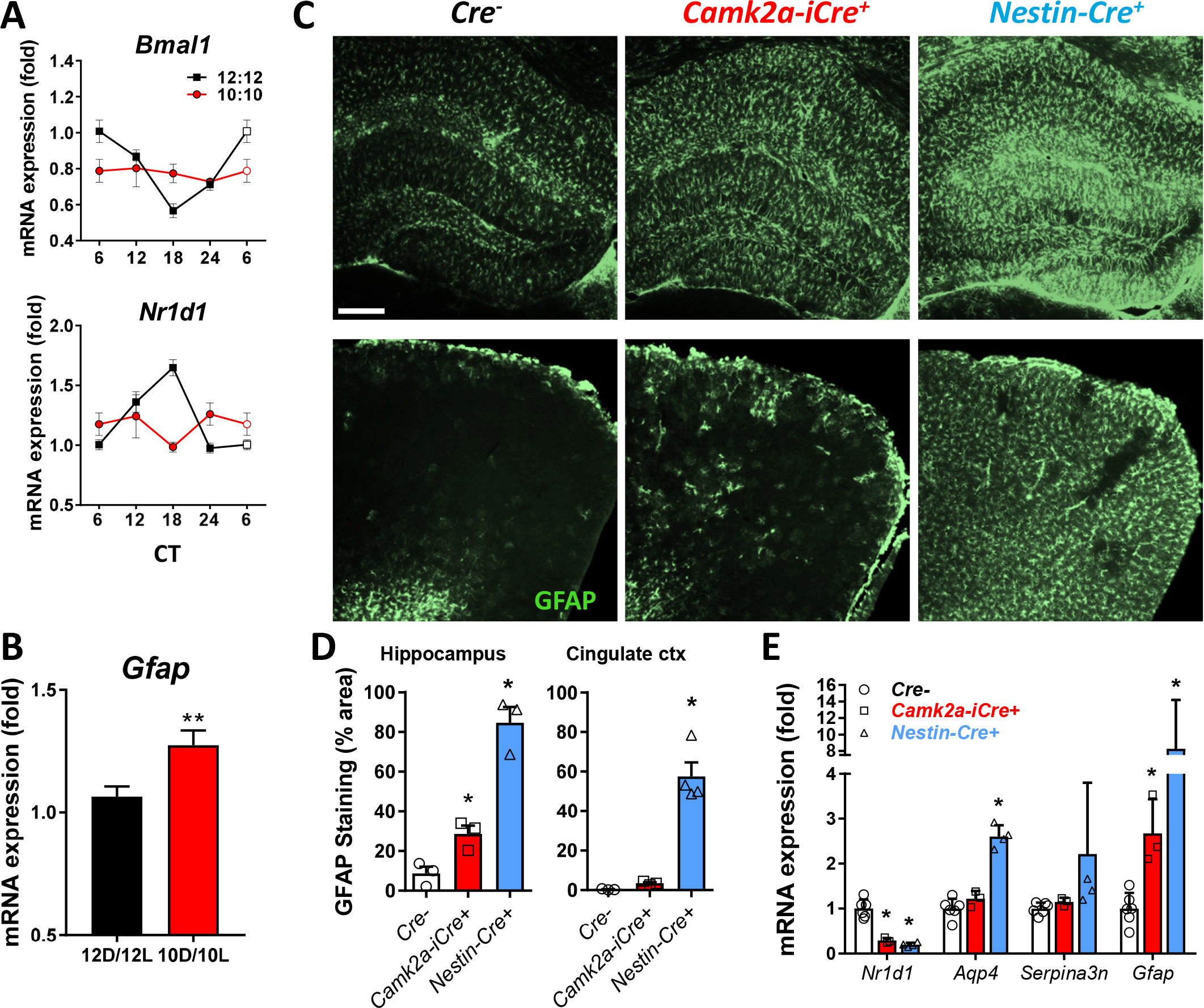
Light-induced circadian disruption or neuron-specific *Bmal1* deletion partially recapitulate the astrocyte activation phenotype observed following brain-specific *Bmal1* deletion. A. Cortical qPCR from 6-week old WT mice housed in 12h:12h or 10h:10h light:dark cycles for 6 weeks, harvested every 6 hours from CT0-CT18 reveals blunting of circadian gene oscillations in 10h:10h mice. B. Total *Gfap* mRNA levels for mice from (A). **p<0.01 by 2-tailed T-test. C. Representative images showing GFAP immunostaining of hippocampus (upper panels) and cingulate cortex (lower panels) of neuron-specific *Bmal1* KO mice (*Cam2a-iCre+;Bmal1^f/f^*) and brain-specific (neurons+astrocytes) *Bmal1* KO mice (*Nestin-Cre+;Bmal1^f/f^*) at 4mo. Scale bar = 100 μm. D. Quantification of GFAP immunoreactivity in two brain regions of Cre-, *Cam2a-iCre+;Bmal1^f/f^*, and *NestinCre+;Bmal1^f/f^* mice. N=3-4/genotype. Each data point=1 mouse. *p<0.05 by 1-way ANOVA. E. Quantification of astrocyte activation transcripts by qPCR in cortex of Cre-;*Bmal1^f/f^*, *Cam2a-iCre+;Bmal1^f/f^*, and *Nestin-Cre+;Bmal1^f/f^* mice. The clock-controlled gene *Nr1d1* is shown to illustrate loss of BMAL1-mediated transcription. *p<0.05 by 2-way ANOVA with Dunnett’s post-test for multiple comparisons. All data represent mean±SEM.

Circadian clock function can be abrogated by deletion of the critical clock gene *Bmal1*. We previously observed that constitutive or post-natal global deletion of *Bmal1* in mice induced striking increases in GFAP+ astrocytes throughout the brain (Musiek et al., 2013; Yang et al., 2016). We asked if the astrocyte activation observed following *Bmal1* deletion was simply a response to neuronal injury, or a cell-autonomous process regulated by the core clock in astrocytes. To address this, we examined the brains of neuron-specific *Bmal1* KO mice (*Camk2a-iCre;Bmal1^f/f^)*. These mice express Cre recombinase in a pan-neuronal manner under the control of a BAC-CaMKIIa promoter, yielding widespread neuronal deletion of *Bmal1* (Fig. S1C) and loss of behavioral circadian rhythms (Izumo et al., 2014). At 4mo, *Camk2aiCre+;Bmal1^f/f^* mice exhibited a modest increase in astrocyte activation, as assessed by GFAP immunostaining (Fig. 1C,D) and qPCR (Fig. 1E), in the cortex and hippocampus. However, similarly-aged *Nestin-Cre+;Bmal1^f/f^* (referred to as NBmal1 KO) mice, in which *Bmal1* is deleted in neurons and astrocytes, have much more prominent astrogliosis throughout the brain and higher expression of activation-related transcripts *Aqp4* and *Serpina3n* (Fig. 1C-E). While NBmal1 KO mice exhibit large increases in GFAP+ astrocytes in the cortex, this did not appear to be due to increased astrocyte division, as NBmal1 KO mice had no significant increase in dividing Ki67+, GFAP+ double-positive cells (Fig. S1D), or in immunoreactivity for the pan-astrocyte marker S100B (Fig. S1E). This finding suggests that while a small component of astrogliosis observed following *Bmal1* deletion may be a response to neuronal injury, BMAL1 within astrocytes appears to play a critical cell-autonomous role in astrocyte activation.

To assess the potential cell-autonomous effects of *Bmal1* on astrocyte activation, we generated primary astrocyte-enriched cultures from *Bmal1* KO mice and WT littermates, and examined activation markers *in vitro*. *Bmal1* KO astrocytes were viable and appeared to have normal morphology, but had significantly elevated expression of *Gfap* and *Aqp4* mRNA and GFAP protein (Fig. 2A,B), suggesting spontaneous activation in the absence of neuronal influence. In order to exclude a developmental influence of BMAL1, we used siRNA to knock down *Bmal1* expression in WT primary astrocyte cultures. This method resulted in a 95% decrease in BMAL1 protein (Fig. 2C,D) and a 66% decrease in expression of the BMAL1 target *Nr1d1* (which encodes REV-ERBα)(Fig. S2A). *Bmal1* knockdown (KD) in astrocytes induced a highly consistent (across 14 independent experiments) 36% increase in *Gfap* mRNA expression (Fig. S2A) as well as a 68% increase in GFAP protein (Fig. 2C,D). *Bmal1* KD in primary astrocytes from *Per2*-luciferase reporter mice caused a loss of circadian *Per2* rhythms (Fig. 2E). We also saw a significant increases in the astrocyte activation marker *S100a6* (1.7 fold), the cytokine *Il6* (2.4 fold), and *Il33* (1.6 fold) (Fig. S2A), an astrocytic cytokine implicated in astrocyte-microglia signaling (Wicher et al., 2017). Thus, disruption of circadian function via knockdown of *Bmal1* in WT astrocytes induces a cell-autonomous reactive phenotype *in vitro*.

**Figure 2.**
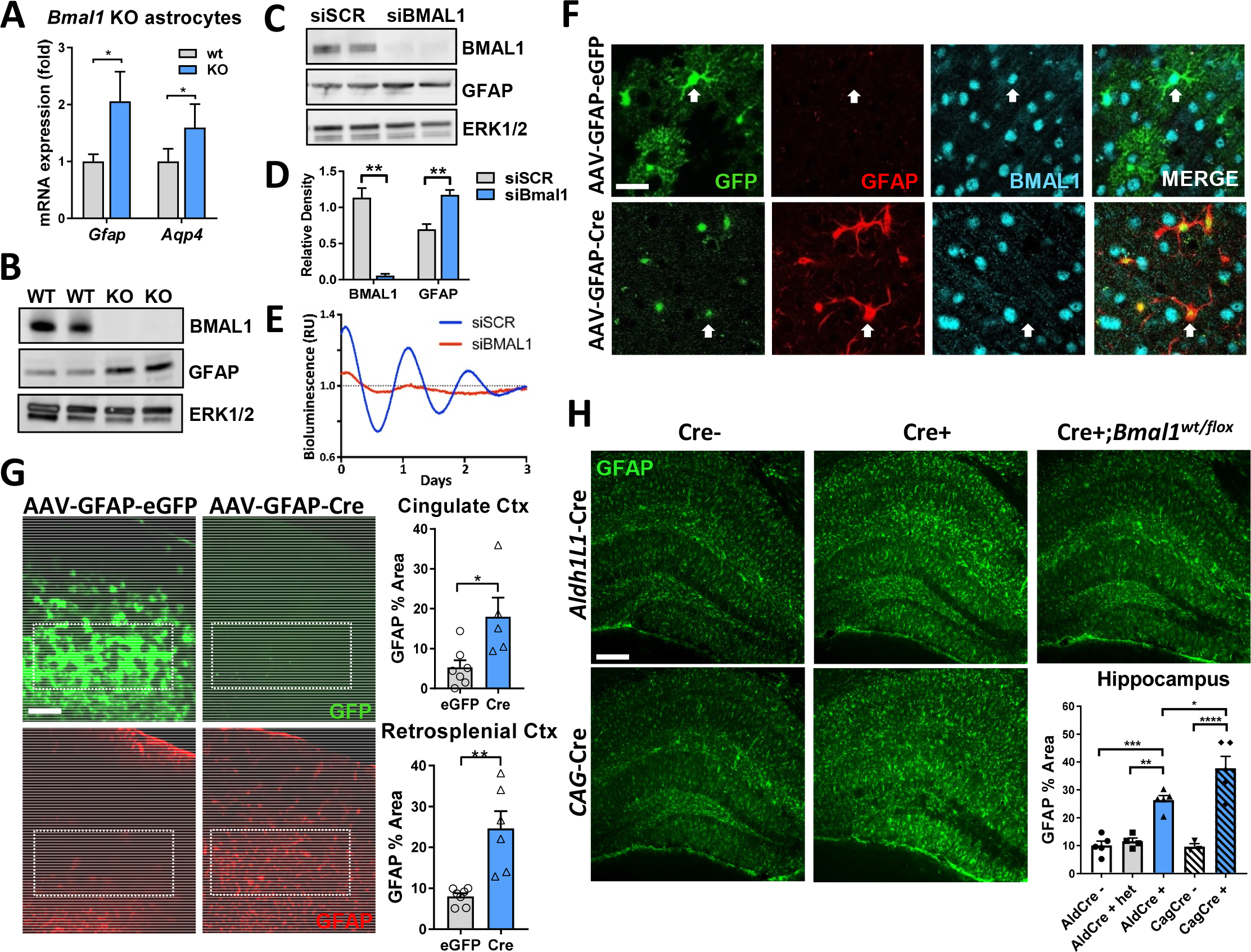
*Bmal1*deficiency induces cell-autonomous astrocyte activation *in vitro* and *in vivo*. A. qPCR of *Gfap* and *Aqp4* mRNA in primary astrocyte-enriched cultures from WT or global *Bmal1* KO mice. N=8-12 samples from 3 independent experiments. B. Western blot showing increased GFAP protein levels in cultured astrocyte cell lysates from *Bmal1* KO mice. In (B) and (C), p42/44 ERK protein is shown as a loading control. C. Western blot showing knockdown of BMAL1 and increased GFAP protein in WT astrocyte cultures 7 days after treatment with non-targeting siRNA (siSCR) or siRNA targeting *Bmal1* (siBmal1). D. Densitometric quantification of blots in (C). N=4 samples/group. E. Representative trace depicting detrended circadian oscillations in luminescence seen in primary astrocyte cultures from *Per2*-Luc reporter mice treated with siRNA (siSCR, blue or siBmal1, red). Similar results were obtained in 3 separate cultures. F. Representative confocal images from cerebral cortex of *Bmal1^f/f^* mice 5 mo after i.c.v. injection at age P2 with viral vectors inducing astrocyte-specific expression of eGFP (AAV8-GFAP-eGFP) or Cre (with a fused eGFP tag, AAV8-GFAP-Cre). AAV8-GFAP-GFP-infected cells show whole-cell GFP expression and persistent colocalized nuclear BMAL1 staining, while AAV8-GFAPCre infected cells show only nuclear GFP expression (indicating the presence of Cre^eGFP^ fusion protein), increased GFAP expression, and loss of nuclear BMAL1. Scale bar= 20µm. G. Representative images (left) and quantification (right) illustrating increased percent area covered by GFAP+ astrocytes in cortex in mice from (F) injected with AAV8-GFAP-Cre, as compared to AAV8-GFAP-GFP-injected controls. Note that AAV8-GFAP-Cre virus encodes a Cre^GFP^ fusion protein which is nuclear-localized and difficult to see in the GFP image, but is present upon higher magnification as in (F). Scale bar= 150µm. Each data point = 1 mouse. H. Representative images and quantification of % area of GFAP staining from hippocampus of *Aldh1l1-Cre^ERT2^+;Bmal1^f/f^* and *Aldh1l1-Cre^ERT2^+;Bmal1^wt/flox^* mice at 3 mo (2 months after tamoxifen treatment), *CAG-Cre^ERT2^+;Bmal1^f/f^* mice at 4 mo (2 months after tamoxifen treatment), and *Bmal1^f/f^* controls. Scale bar = 150µm. All data represent mean+SEM. *p<0.05, **p<0.01, ***p<0.001, ****p<0.0001 by 2-tailed T-tests (C) and (G) or 1-way ANOVA (H) with Holm-Sidak correction for multiple comparisons when applicable.

We next sought to confirm these findings by specifically deleting *Bmal1* in astrocytes *in vivo*. We found that intracerebroventricular injection of an AAV8 viral vector expressing eGFP under a *Gfap* promoter (AAV8-GFAP-eGFP) on postnatal day 2 led to widespread eGFP expression in resting astrocytes throughout the cerebral cortex, with almost no neuronal labeling (Fig. 2F,G, S2B). Thus, we injected P2 *Bmal1^f/f^* mice either with AAV8-GFAP-GFP, or with an identical virus expressing a Cre-GFP fusion protein (AAV8-GFAP-Cre^GFP^). At 5mo, we observed *Bmal1* deletion and increased GFAP expression specifically in astrocytes expressing AAV8-GFAP-Cre^GFP^ (Fig. 2F). It should be noted that the fusion product of the GFAP-Cre^GFP^ vector exhibits very weak nuclei-restricted fluorescence, making it much more difficult to see than that of the AAV8-GFAP-eGFP vector. Mice treated with the eGFP control vector had widespread astrocyte labeling throughout the cingulate and retrosplenial cortices, intact BMAL1 expression, and very little astrocyte activation as defined by GFAP immunoreactivity (Fig. 2F,G). Mice injected with the Cre^GFP^ vector showed astrocytic nuclear GFP expression and 3.4- and 3.1-fold increases in GFAP immunoreactivity in the cingulate and retrosplenial cortices, respectively (Fig. 2G). Finally, we generated tamoxifen-inducible astrocyte-specific *Bmal1* KO mice using an *Aldh1l1-*Cre^ERT2^ driver line (*Aldh1l1*-Cre^ERT2^;*Bmal1*^f/f^ mice) (Srinivasan et al., 2016). We treated Cre+ and Cre- mice with tamoxifen at 1mo, and examined astrocyte activation at 3mo. We observed specific deletion of *Bmal1* in astrocyte nuclei of the hippocampus and cortex (Fig. S2D,E). *Aldh1l1*-Cre+ mice had significant increases in astrogliosis, as assessed by GFAP immunostaining in the hippocampus (Fig. 2H) and a nearly significant (p=0.055) increase in the retrosplenial cortex (Fig. S2C). While it is difficult to directly compare tamoxifen-inducible and constitutive Cre lines, *Aldh1l1*-Cre+ mice had similar levels of astrogliosis to tamoxifen inducible global *Bmal1* KO mice (*CAG-Cre^ERT2^+;Bmal1^f/f^*)(Yang et al., 2016) also harvested 2mo after TAM treatment (Fig. 2H, S2C). Astrogliosis was not caused by Cre expression, as Cre+;*Bmal1^flox/wt^* mice had no phenotype (Fig. 2H, S2C). *Ald1l1-Cre+* mice also had spontaneous and significant increases in a number of additional astrocyte activation marker transcripts including *S100a6* (2.5 fold), *C4b* (2.6 fold), *Cxcl5* (7.5 fold), and *Mmp14* (3.1 fold) (Fig. S2F). In total, these experiments demonstrate that *Bmal1* deletion in astrocytes can induce activation in a cell-autonomous manner.

While activated astrocytes generally exhibit morphologic changes and increased GFAP expression, recent studies have demonstrated that the transcriptional profile of activated astrocytes is heterogeneous and can define their function (Liddelow et al., 2017; Zamanian et al., 2012). In order to examine the transcriptional profile in the brain following *Bmal1* deletion, we performed microarray analysis of cortical tissue from 11mo NBmal1 KO mice, Cre-controls, and *Per1*^Brdm^/*Per2*^Brdm^ (*Per1/2*^mut^) mice, placed in constant dark conditions for 36 hours then harvested at CT 6:00 and 18:00. *Per1/2*^mut^ mice were included because they are behaviorally arrhythmic and have a dysfunctional circadian clock, but have no astrogliosis phenotype as well as tonically increased levels of BMAL1 transcriptional targets (Fig. 3A, S3A). To understand astrocyte transcriptional changes in NBmal1 KO brain, we examined the overlap between the 200 most upregulated transcripts in NBmal1 KO, with the 200 most astrocyte-specific transcripts, according to the Barres Lab RNAseq database (Zhang et al., 2014b) (Fig. 3A). Thirty genes overlapped, including *Gfap, Aqp4* (encoding the astrocytic water channel Aquaporin-4), and *Megf10*, a gene involved in astrocyte-mediated synaptic elimination (Chung et al., 2013). *C4b*, a complement cascade component also implicated in synapse elimination and a marker of astrocyte activation and aging (Boisvert et al., 2018; Clarke et al., 2018), was strongly upregulated (4-fold) in NBmal1 KO brain. Several other disease-associated astrocytic genes were increased in NBmal1 KO brain, including inflammatory mediators such as *Cxcl5* (Bortell et al., 2017) and *Ccr7* (Gomez-Nicola et al., 2010), calcium binding protein genes *S100a4* (Dmytriyeva et al., 2012) and *S100a6* (Hoyaux et al., 2002), and matrix metaloproteinases *Mmp14* (Rathke-Hartlieb et al., 2000) and *Mmp2* (Li et al., 2011). Nearly all of these transcripts were either unchanged or decreased in *Per1/2*^mut^ mice, demonstrating the key role of decreased BMAL1 function in this phenotype. We next performed microfluidic qPCR profiling of global *Bmal1* KO hippocampal samples for markers of astrocyte activation and polarization (Liddelow et al., 2017), which demonstrated that most pan-reactive markers were increased, while *Bmal1* deletion did not clearly polarize astrocytes toward an neurotoxic or neurotrophic profile (Fig. 3B). Taken together, these genetic profiling studies suggest that *Bmal1* deletion induces a unique state of astrocyte activation with potential implications for astrocyte behavior in disease states.

**Figure 3.**
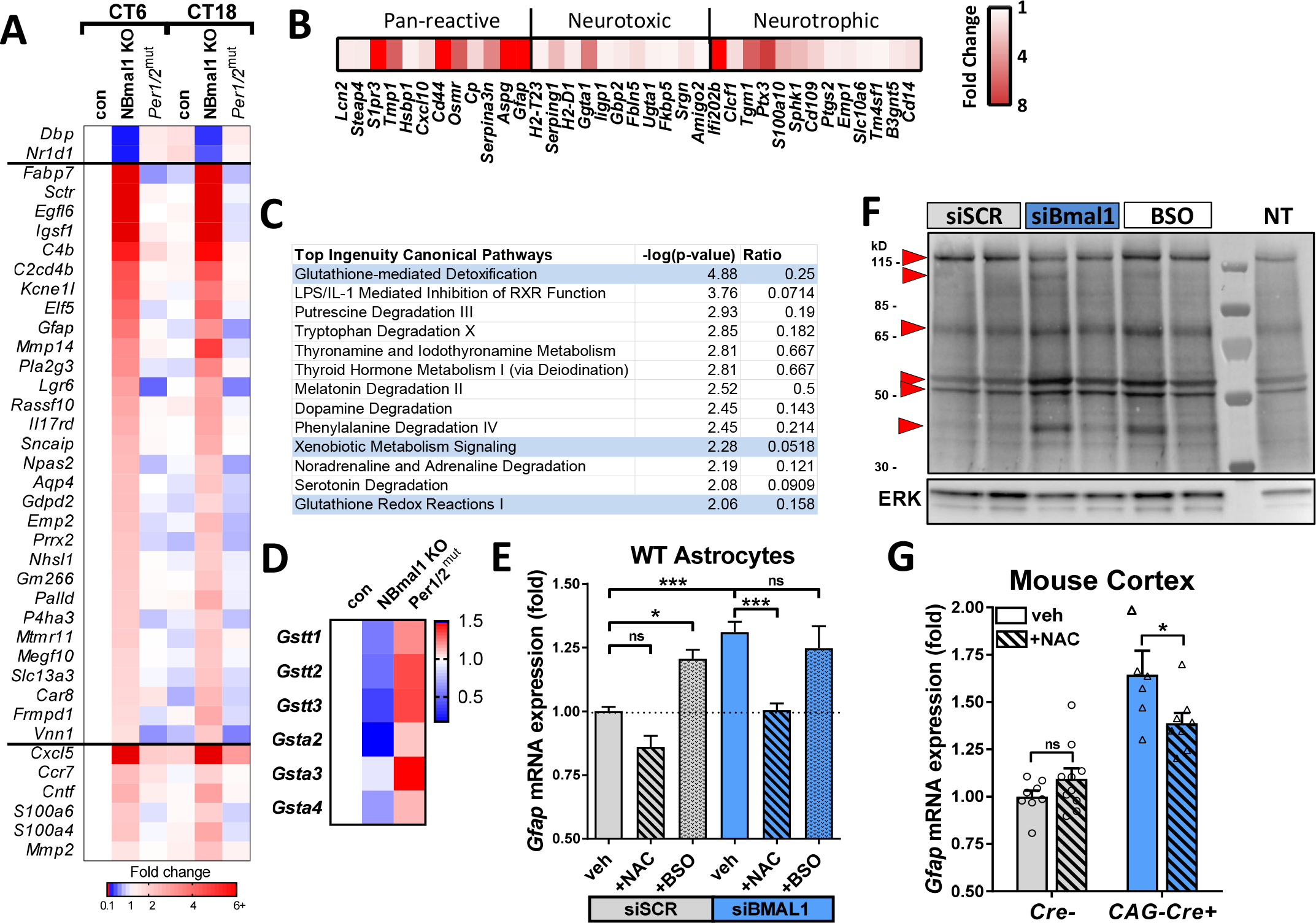
BMAL1 regulates astrocyte activation via a glutathionylation-dependent mechanism. A. Astrocyte-specific genes upregulated in *Nestin*-Cre^+^;Bmal1^f/f^ (NBmal1 KO) cortex and changes to the same genes in *Per1/2^mut^* mice. Transcripts upregulated at least 2-fold in NBmal1 KO were compared to the top 200 astrocyte-specific genes (Zhang et al, 2014), and overlapping genes are displayed. Bottom - other notable genes implicated in astrocyte activation and function which were upregulated in NBmal1 KO cortex. The clock-controlled genes *Dbp and Nr1d1* are direct *Bmal1* transcriptional targets. B. Expression of neurotoxic, neurotrophic, and pan-reactive astrocyte activation genes (Liddelow et al., 2017) in 5 mo global *Bmal1* KO vs WT littermate hippocampus assayed by microfluidic qPCR. Each datapoint represents the average of 4 *Bmal1* KO mice normalized to the average of 4 WT littermates. C. Results from Ingenuity Pathway Analysis Canonical Pathway Analysis of NBmal1 KO microarray data. Pathways related to glutathione homeostasis are colored blue. D. Expression levels for downregulated glutathione-S-transferases in NBmal1 KO cortex vs. *Bmal1^f/f^* controls. Expression levels in *Per1/2^mut^* cortex included for comparison. Based on microarray data. E. qPCR illustrating effect of glutathione manipulation on *Gfap* transcript levels following *Bmal1* knockdown. WT primary astrocytes treated with control siRNA (siSCR, grey bars) or siBmal1 (blue bars), +/- N-acetyl-cysteine (NAC, striped bars) to increase glutathione levels, or buthionine sulfoxime (BSO, speckled bars) to deplete glutathione. N=9 samples/condition from 3 separate experiments. F. WT astrocytes transfected with siSCR, siBmal1, or siSCR +BSO (BSO) were treated with biotin-linked glutathione ethyl ester (bioGEE) to assess glutathionylation. Increased band intensity indicates *decreased* endogenous glutathionylation. Arrows denote altered bands. NT = not treated with bioGEE, showing non-specific labeling. G. qPCR depicting effects of NAC treatment on *Gfap* transcript levels of inducible *Bmal1* KO mice (*CAGCre^ERT2^+;Bmal1^f/f^*). Inducible *Bmal1* KO mice and Cre- control littermates were treated with NAC prior, during, and after 5-day tamoxifen treatment to delete *Bmal1*. Mice were harvested 9 days after start of tamoxifen, and *Gfap* levels were quantified in cortex. N=6-10 mice per group. All data represent mean+SEM. *p<0.05, ***p<0.001 by T-test with Holm-Sidak correction for multiple comparisons.

Although *Per1/2^mut^* and NBmal1 KO mice are both behaviorally arrhythmic, expression of BMAL1 transcriptional targets *Nr1d1* and *Dbp* are elevated solely in the *Per1/2^mut^* mice, due to loss of inhibition of BMAL1 (Fig. 3A, S3A). *Gfap* is not elevated *Per1/2^mut^* mice (Fig. 3A, S3A), suggesting that loss of BMAL1 activity, or at least blunting of BMAL1 oscillation (as in Fig. 1A), is needed to induce astrocyte activation. In order to determine if *Gfap* and other selected activation markers are, in fact, clock controlled genes, we harvested cortex from 4mo NBmal1 KO and Cre- controls at 4-hour intervals across one circadian cycle in constant darkness. *Gfap* mRNA levels showed no circadian oscillation in control or Cre+ mice, and were elevated across the circadian cycle in the cortex of Cre+ mice. These findings were duplicated in other markers of astrocyte activation, including *Aqp4* and *Mmp14* (Fig. S3B). The lack of circadian oscillation suggests that this astrogliosis phenotype is not due to direct regulation of *Gfap* and other activation markers by the clock.

To address potential mechanisms linking BMAL1 to astrocyte activation, we performed pathway analysis on our NBmal1 KO transcriptomic data using two bioinformatics tools: Ingenuity Pathway Analysis (IPA) and DAVID Bioinformatics Resources 6.8. We analyzed an inclusive gene list consisting of all genes that differed between control and NBmal1 KO at 6pm with uncorrected p values<0.01 (Table S1). IPA Canonical Pathway Analysis identified Glutathione-mediated detoxification as the top hit (Fig. 3C, p= 0.000013). Functional Annotation Clustering with DAVID showed that the most enriched gene cluster focused on glutathione-s-transferase (GST) activity, and 4 of the top 5 KEGG pathways identified were related to glutathione metabolism (Fig. S3C). GST enzymes catalyze the phase II detoxification of reactive intermediates in cells via adduction to glutathione and can alter cellular signaling via glutathionylation of proteins (Grek et al., 2013). We found that expression of numerous GST isoforms was suppressed in NBmal1 KO cortex, and were increased in *Per1/2*^mut^ cortex (Fig. 3D), suggesting reciprocal regulation by the positive and negative limbs of the circadian clock. In support of our microarray analysis, we observed significant decreases in 2 of the 3 GST isoforms probed in cortex from 3mo *CAG-Cre+;Bmal1^f/f^* mice 9 days after tamoxifen treatment (Fig. S3D) and in 1 of the 3 GST isoforms probed in hippocampus from 3mo *Aldh1l1-Cre+* mice 2 months after tamoxifen treatment (Fig. S3E). We next asked if altered glutathione homeostasis within astrocytes linked *Bmal1* deficiency with astrocyte activation. Treatment with N-acetyl cysteine (NAC), a glutathione precursor, completely prevented increased *Gfap* expression in astrocytes following *Bmal1* knockdown (Fig. 3E). Conversely, incubation of WT astrocytes with buthionine sulfoxime (BSO), which blocks cysteine uptake and depletes cellular glutathione levels, lowered cellular glutathione levels (Fig. S3F) and caused a similar increase in *Gfap* mRNA to that observed with *Bmal1* knockdown (Fig. 3E). Treatment of *Bmal1* deficient astrocytes with BSO did not induce a further increase in *Gfap* suggesting that BSO and *Bmal1* knockdown are causing increases in *Gfap* through overlapping pathways. Interestingly, neither *Bmal1* KO nor knockdown of *Bmal1* in cultured primary astrocytes altered glutathione levels (Fig. S3F). Additionally, the oxidation state of glutathione (ratio of reduced GSH to oxidized GSSG) was not altered in cortex from NBmal1 KO or *Per1/2^mut^* mice (Fig. S3G). As glutathione levels and oxidation state remained unchanged with loss of *Bmal1*, we next examined glutathione signaling via glutathionylation, which has been shown to oscillate in a circadian manner in the SCN (Wang et al., 2012). Western blot analysis of siSCR and siBmal1 transfected WT astrocytes treated with biotin-linked glutathione ethyl ester (bioGEE) revealed widespread changes in the pattern of s-glutathionylated proteins in *Bmal1* KD cells, which are similar to changes seen in BSO treated controls (Fig. 3F). In general, numerous bands showed increased biotin labeling, indicating decreased glutathionylation, as would be expected with decreased GST expression. These data suggest that the observed astrogliosis may be due at least in part to altered protein glutathionylation resulting from impaired expression of specific GSTs.

We next examined whether bolstering glutathione signaling can prevent astrogliosis following *Bmal1* deletion *in vivo*. We previously showed that tamoxifen-inducible global *Bmal1* deletion in mice (*CAG-Cre+;Bmal1^f/f^* mice) causes astrocyte activation (Yang et al., 2016). We treated *CAG-Cre+;Bmal1^f/f^* mice with NAC in their drinking water (40mM beginning 5 days before tamoxifen) and via intraperitoneal injection (150mg/kg, daily beginning 2 days before tamoxifen), and induced *Bmal1* deletion via tamoxifen administration for 5 days. Both administrations of NAC were continued for the duration of the experiment. Cortex was harvested 9 days after the start of tamoxifen treatment and analyzed by qPCR. Tamoxifen treatment induced an 87% loss in *Bmal1* and a 79% loss in expression of the *Bmal1* target *Nr1d1* (Fig. S3H), as well as a 64% increase if *Gfap* expression (Fig. 3G). Administration of NAC was able to provide a significant, but partial rescue of the *Gfap* increase seen in *CAG-Cre+;Bmal1^f/f^* animals (Fig. 3G). These results suggest that *Bmal1* regulates *Gfap* expression at least in part through modulation of glutathione signaling *in vivo*.

Finally, we examined potential pathogenic consequences of *Bmal1* deletion in astrocytes. We generated primary astrocyte-enriched cultures from *CAG-Cre+;Bmal1^f/f^* mice and Crelittermates and treated them with tamoxifen *in vitro* after culturing to delete *Bmal1*. These cultures are primarily astrocytes, but this methods yields cultures which contain ~2% microglia and/or oligodendrocyte precursor cells (Schildge et al., 2013). Primary WT cortical neurons were then co-cultured on these astrocytes for up to 12 days and a subset were subjected to oxidative stress by hydrogen peroxide exposure (100µM H_2_O_2_) for 24 hours (Fig. 4A). We found no difference in the number of neurons surviving on Cre- and Cre+ astrocyte cultures 1 day after plating (as quantified by staining for neuronal nuclei with NeuN, as these cells do not yet have many MAP2 positive neurites) (Fig. 4B). At day 7 after plating, there was a 45% decrease in the number of NeuN+ nuclei (Fig. 4B,C), a trend toward a decrease in MAP2 staining, and a trend toward an increase in cleaved-caspase-3 staining (Fig. S4A,B) in the Cre+ astrocyte condition. The decrease in MAP2 staining became significant after 12 days (Fig. 4D,E). H_2_O_2_ exposure at day 12 decreased MAP2 levels in Cre- cultures, and the combination of *Bmal1* deletion and H_2_O_2_ killed nearly all neurons (Fig. 4D,E). This finding shows that *Bmal1*-deficient astrocytes are less able to sustain neuronal viability, particularly in the face of oxidative stress.

**Figure 4.**
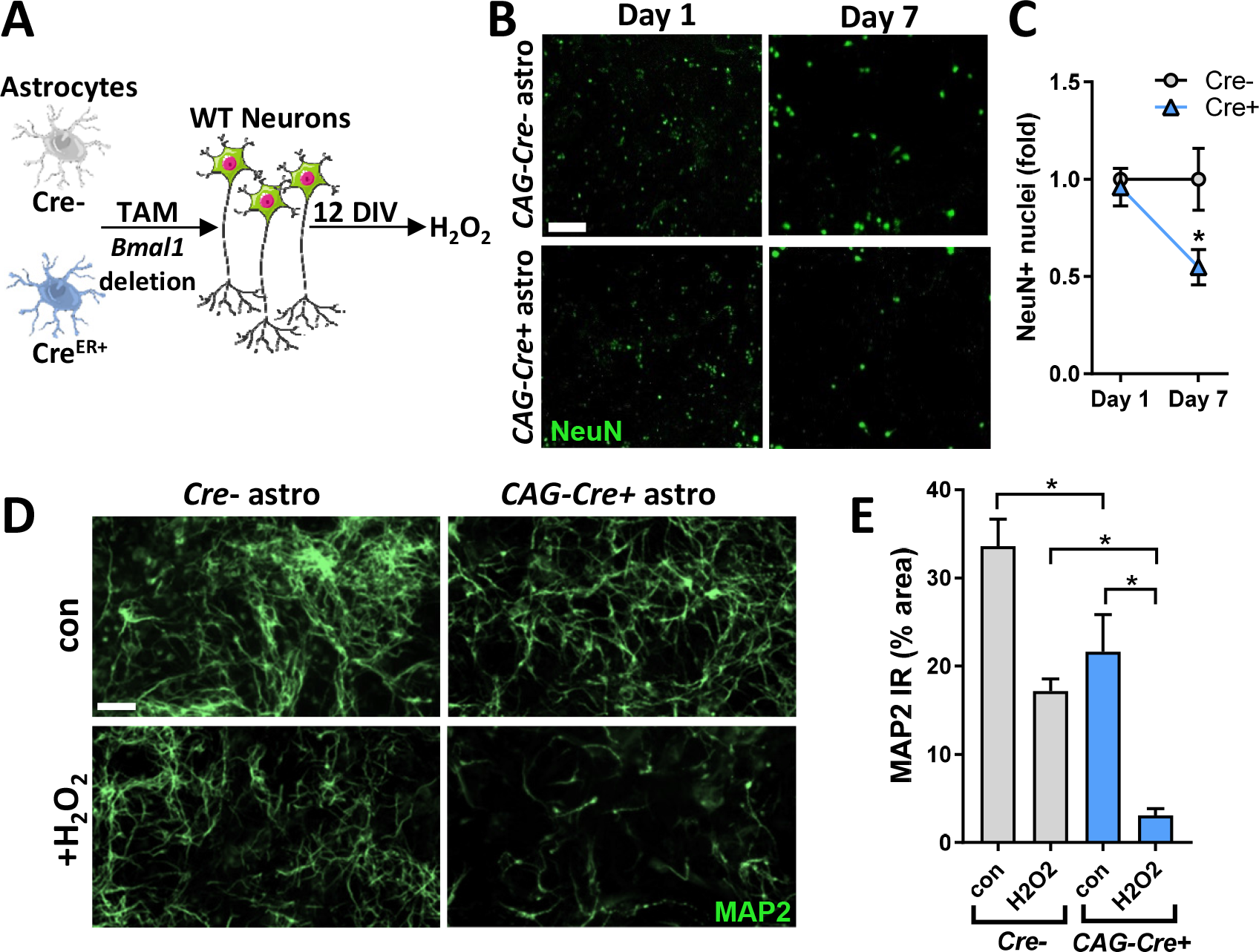
Loss of *Bmal1* impairs astrocytic support of neurons and astrocytic amyloid plaque clustering. A. Representative images showing NeuN+ WT neuronal nuclei at DIV 1 and 7, grown on primary astrocytes from *Cre-*control mice, or inducible *Bmal1* KO mice (*iCAG-Cre+*), treated with tamoxifen *in vitro*. Scale bar = 50μm. B. Quantification of NeuN+ nuclei from (A) normalized to NeuN counts from Cre- condition. N=8 replicates from 2 independent experiments. C. Representative images showing MAP2+ WT neurons grown as in (A) at DIV 12. Lower panels show neurons from adjacent wells treated for 24 hours with hydrogen peroxide at DIV 11 (H_2_O_2_, 100µM). Scale bar = 50μm. D. Quantification of MAP2 immunoreactivity (% area) from (C). N=9 replicates from 3 independent experiments. All data are represented as mean±SEM. *p<0.05, **p<0.01 by 2-tailed T-test.

## Discussion

Despite a large body of evidence describing circadian disruptions in both aging and neurodegenerative disease, little is known about the implications of circadian dysfunction on the brain. Here, we provide evidence that behavioral circadian disruption or disruption of the astrocytic molecular clock via manipulations of the master clock gene *Bmal1* induce astrogliosis and astrocyte dysfunction. The critical, but complex role of astrocytes in brain pathologies and neuroinflammation underscores the importance of elucidating both triggers and mechanisms of astrocyte activation, and our data illuminate BMAL1 as a potent regulator of astrocyte activation by both cell-autonomous and non-autonomous mechanisms.

The existence of the astrocyte circadian clock has been well documented (Prolo et al., 2005), and the critical role of the astrocyte clock in maintaining circadian rhythms in the SCN has recently been described (Barca-Mayo et al., 2017; Brancaccio et al., 2017; Tso et al., 2017). Clock genes have been shown to regulate astrocytic glutamate uptake and ATP release (Beaule et al., 2009; Marpegan et al., 2011), but the function of the core clock in astrocytes is otherwise unexplored. We have shown here, using several *in vitro* and *in vivo* methods, that loss of *Bmal1* function in astrocytes causes cell-autonomous astrocyte activation. However, deletion of *Bmal1* specifically in neurons also induces a partial astrocyte activation phenotype, suggesting a coexistent cell non-autonomous mechanism. We have previously shown that *Bmal1* deletion in cultured neurons causes toxicity (Musiek et al., 2013), implicating a possible damage signal from *Bmal1*-deficient neurons which activates nearby astrocytes. Alternatively, the neuron-specific *Bmal1* KO mice used in this study are behaviorally arrhythmic (Izumo et al., 2014), and this loss of rhythms can induce astrocyte activation, as seen in 10h:10h L:D exposed animals (Fig. 1). Thus, cell non-autonomous influences on glial activation in the brain must be examined in the future.

One important question is whether the loss of rhythmic function of the circadian clock plays a key role in this phenotype, or if this is a “non-circadian” function of BMAL1. Our data suggest that arrhythmicity in the setting of increased BMAL1 expression, as in *Per1/2^mut^* mice, does not induce astrocyte activation, as these mice express increased levels of GSTs. Thus, astrocyte activation appears to be dependent on suppression of BMAL1-mediated transcription. However, BMAL1 DNA binding is highly rhythmic and regulated by the clock (Koike et al., 2012), making it difficult to separate entirely from circadian rhythms. *Bmal1* knockdown in astrocytes abrogates circadian *Per2*-luc oscillations, which could contribute to increased *Gfap* expression in cell culture. Moreover, we show in Fig. 1 that circadian disruption by exposure of mice to 10h:10h L:D conditions, which blunts *Bmal1* oscillations in the brain, can induce *Gfap* expression. Thus, dampening the rhythmicity of BMAL1 expression through circadian clock disruption likely plays a role in mediating astrogliosis, perhaps by restricting BMAL1 levels at certain key times of day. BMAL1 levels can be suppressed or blunted by several factors related to neurodegeneration, such as aging (Wyse and Coogan, 2010), inflammation (Curtis et al., 2015), or amyloid-beta (Song et al., 2015), all of which could potentially influence astrocyte activation and function in a BMAL1-dependent manner.

The activation profile induced in *Bmal1*-deficient astrocytes encompasses upregulation of a variety of genes across the pre-defined neurotoxic, neurotrophic, and pan-reactive categories (Liddelow et al., 2017). However, it is important to remember that these astrocyte phenotypes are thought to be induced by cytokine release from microglia (Liddelow et al., 2017), whereas activation due to loss of *Bmal1* can be cell-autonomous. Our findings contrast with those of Nakazato et al, who found that *S100b*-Cre+;*Bmal1*^f/f^ mice did not develop astrocyte activation, and attributed the astrocyte activation phenotype of *Nestin*-Cre+;*Bmal1*^f/f^ mice to pericyte dysfunction (Nakazato et al., 2017). However, the degree of astrocyte *Bmal1* deletion in the *S100b*-Cre mouse was not demonstrated in that study. While our data does not exclude a contribution of blood-brain barrier dysfunction to the *Nestin-Bmal1* phenotype, we provide multiple lines of evidence demonstrating cell-autonomous astrocyte activation. While *Bmal1-*deficient astrocytes do not assume a clear neurotoxic (“A1”) phenotype, our data demonstrate that they express many activation and pro-inflammatory markers, and that they are less able to support neuronal survival in a co-culture system. In addition to *Gfap*, *C4b*, a complement component and marker of astrocyte activation and aging (Boisvert et al., 2018; Clarke et al., 2018), is strongly increased following *Bmal1* deletion, as are several other pan-reactive markers including *Timp1* and *Serpina3n* (Liddelow et al., 2017). Increased expression of several genes implicated in neuroinflammation and neurodegeneration, including *Megf10* (Chung et al., 2013; Iram et al., 2016), which mediates astrocytic synapse elimination, and *Pla2g3*, a phospholipase implicated in ROS-induced neuronal damage (Martinez-Garcia et al., 2010) was observed. Activation marker *Lcn2* (Lipocalin 2), an anti-inflammatory mediator in the brain (Kang et al., 2017), was not increased, again suggesting a pro-inflammatory state.

Our data support a mechanism in which loss of BMAL1 induces astrogliosis through disruption of GST-mediated protein glutathionylation. In addition to its role in quenching oxidative stress, glutathione can form adducts on cysteine residues in a process termed glutathionylation, which is catalyzed by glutathione-s-transferases (Grek et al., 2013). Glutathionylation can serve as a signaling mechanism in processes such as mitochondrial metabolism (Mailloux and Treberg, 2016) and apoptosis (Naoi et al., 2008). Indeed, circadian rhythms of protein s-glutathionylation in the SCN may regulate neuronal excitability (Wang et al., 2012). Because *Bmal1*-deficient astrocytes have normal total glutathione levels, it is likely that disruption of GST-mediated glutathione signaling or utilization, not glutathione depletion, mediates BMAL1-controlled astrogliosis. Accordingly, we observed an altered pattern of protein s-glutathionylation in *Bmal1-*deficient astrocytes, which resembled that seen in BSO-treated cells. Thus, supplementing *Bmal1*–deficient astrocyte cultures or mice with NAC presumably prevents astrogliosis by promoting non-enzymatic glutathionylation of protein targets, thereby circumventing the loss of GSTs (Grek et al., 2013). Further studies are needed to identify specific glutathionylation targets that mediate astrocyte activation, and to understand regulation of astrocyte glutathione signaling in health and disease.

In summary, our data identify the circadian clock protein BMAL1 as a unique, cell-autonomous regulator of astrogliosis and posit altered GST-mediated protein glutathionylation as a mediator of this effect. Our findings also show that non-genetic circadian disruption promotes *Gfap* expression, and demonstrate that *Bmal1*-deficient astrocytes have diminished neurotrophic function and impaired engagement of amyloid plaques. BMAL1 serves as a novel link between the core circadian clock and astrogliosis, and this study provides new insights into how the circadian clock might influence neurodegeneration.

## Acknowledgements

The authors declare no financial conflicts of interest. This research was funding by NINDS grant K08NS079405 (ESM), NIA grant R01AG054517 (ESM), and a New Investigator Research Grant from the Alzheimer’s Association (ESM). JST is an Investigator in the Howard Hughes Medical Institute. SAL was supported by a postdoctoral fellowship from the Australian National Health and Medical Research Council (GNT1052961), the Glenn Foundation Glenn Award, an anonymous donation, and the generous support of V. and S. Coates.

---

**Supplemental Figure 1:**
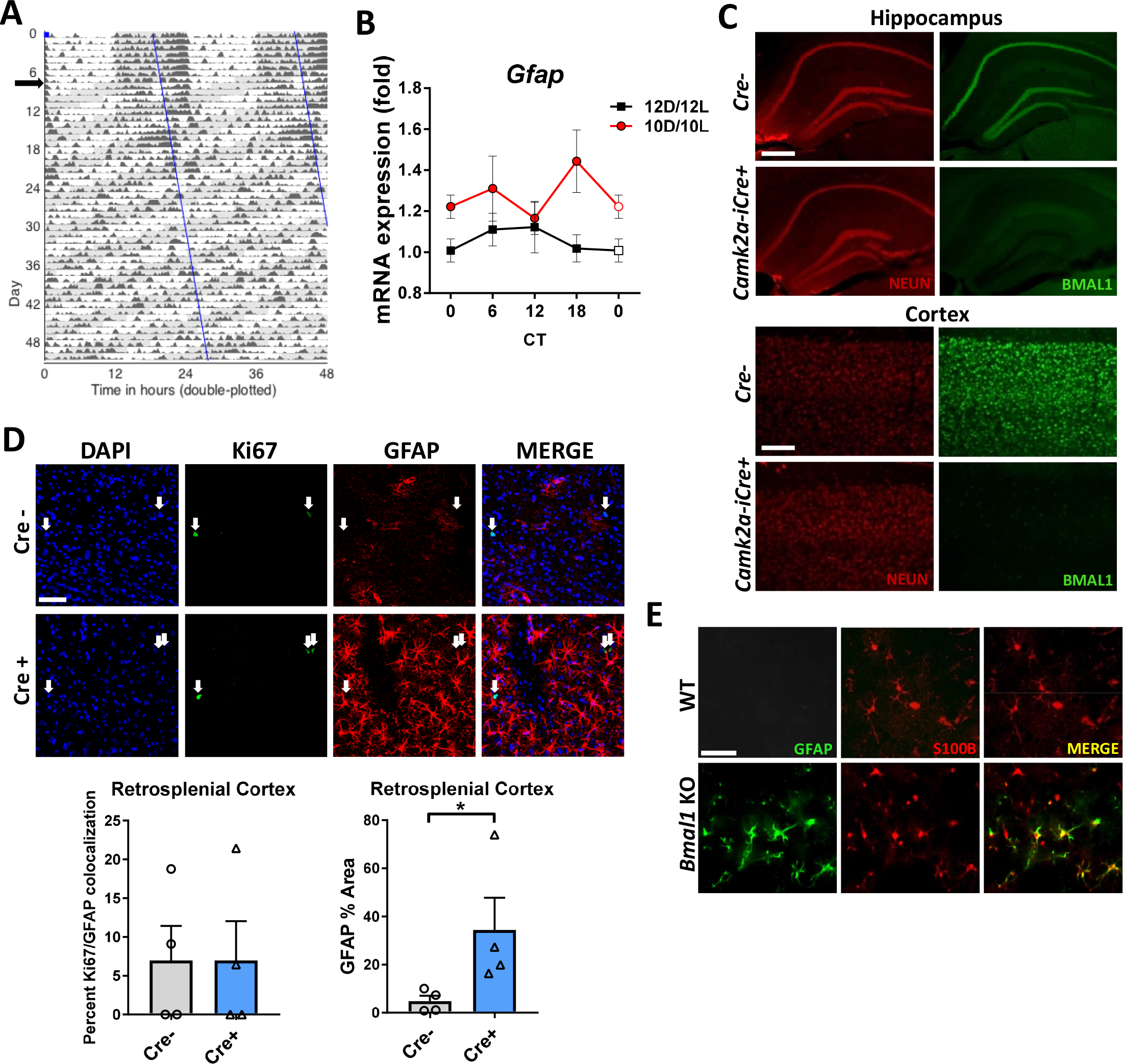
Characterization of mouse models with astrogliosis induced by loss of *Bmal1*. A. Representative actigraph of a WT mouse housed in 12h:12h light:dark, then switched to 10h:10h light:dark at black arrow, leading to behavioral arrhythmicity. B. Cortical qPCR showing mRNA levels from 10h:10h light:dark WT mice from Fig. 1A, S1A reveals increased *Gfap* expression at most timepoints throughout the circadian day. C. Hippocampal (top, scale bar = 400μm) and cortical (bottom, scale bar = 150μm) staining of NEUN (red) and BMAL1 (green) from 4mo *Camk2a-iCre+;Bmal^f/f^* mice and Cre- controls shows complete loss of neuronal BMAL1 in Cre+ animals. D. Representative images (top) and quantification (bottom) illustrating minimal localization of the cell division marker Ki67 staining with GFAP+ astrocytes in cerebral cortex from 4 mo *Nestin-Cre+;Bmal1^f/f^* (NBmal1 KO) mice (left graph), despite large increases in GFAP+ cells as compared to Cre- controls (right graph). Scale bar= 50μm. E. NBmal1 KO mice have increased GFAP+ activated astrocytes in the cerebral cortex without major changes in number of total (S100B+) astrocytes. Scale bar = 50μm. All data represent mean+SEM. *p<0.05 by 2-tailed T-test.

**Supplemental Figure 2.**
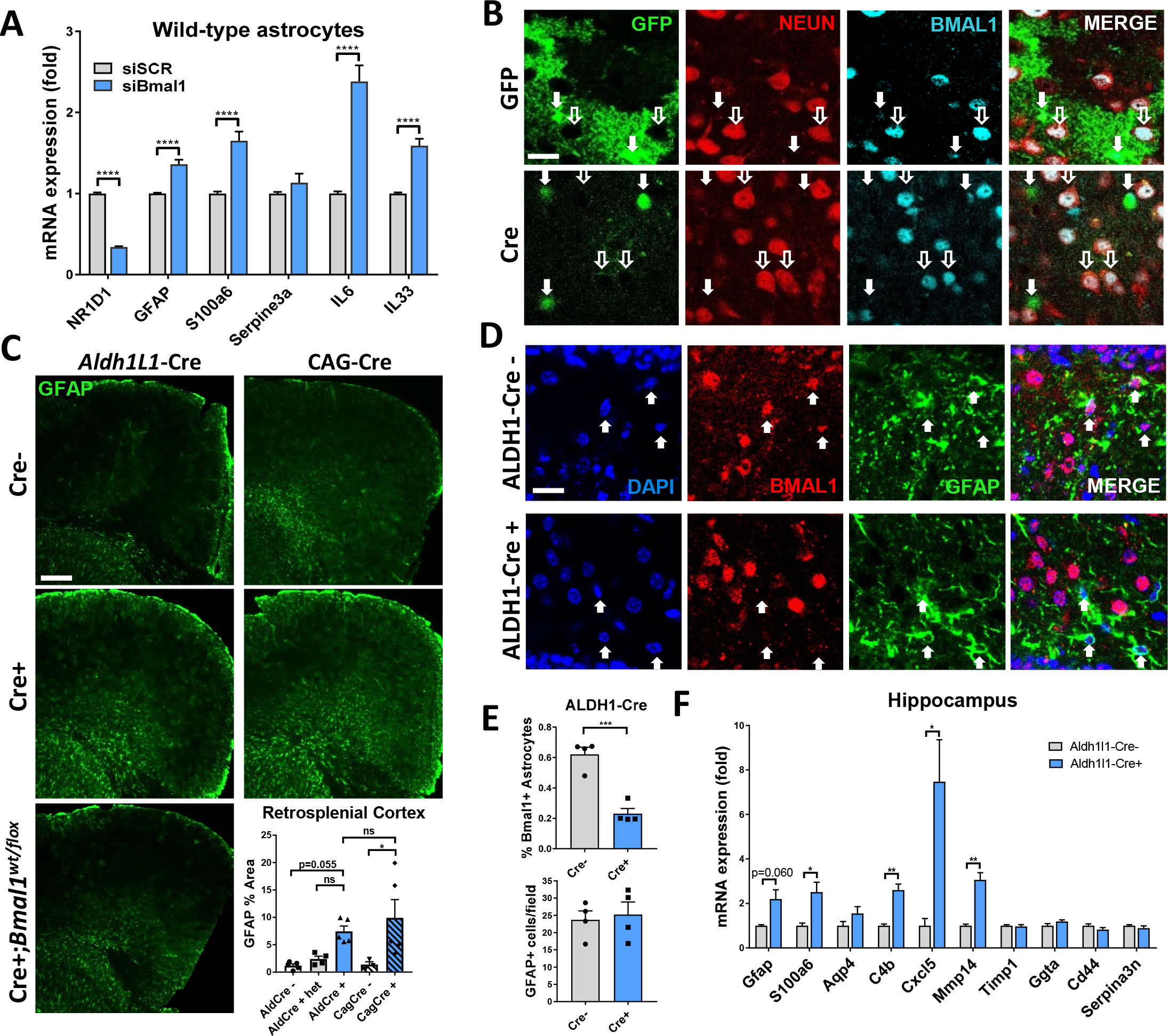
Characterization of astrogliosis seen with loss of *Bmal1* in *vitro* and *in vivo*. A. qPCR showing mRNA levels of several astrocyte activation and inflammatory genes 7 days after treatment of WT astrocytes with non-targeting siRNA (siSCR) or siRNA targeting *Bmal1* (siBmal1). N=12-15 experiments. B. Representative cortical images from mice in Fig. 2F-H. *Top row*. AAV8-GFAP-GFP-infected cells (solid arrows) show whole-cell GFP expression, persistent colocalized nuclear BMAL1, and fail to colocalize with neuronal nuclei (hollow arrows). *Bottom row*. AAV8-GFAP-Cre infected cells show only nuclear GFP expression (solid arrows, Cre^eGFP^ fusion protein), loss of nuclear BMAL1 and fail to colocalize with neuronal nuclei (hollow arrows). Scale bar= 20µm. C. Representative images and quantification of percent GFAP area from retrosplenial cortex of mice in Fig. 2H. Scale bar = 150µm. D. Representative confocal images of *Aldh1l1-Cre+;Bmal^f/f^* mice and Cre- controls. DAPI+, GFAP+ (solid arrows) astrocytes colocalize with BMAL1 in Cre- mice, but not in Cre+ mice. Scale bar= 40µm. E. Quantification of DAPI/GFAP/BMAL1+ cells (top) or GFAP+ cells per field of view (bottom) in *Aldh1l1-Cre+;Bmal^f/f^* mice vs Cre- controls in hippocampus. Each data point = 1 mouse. 6 fields/mouse with an average of 24.5 GFAP+ cells/field, totaling 1,176 cells counted. Note that there is not an increase in GFAP+ cells in hippocampus, just the size and shape of these cells. F. mRNA levels of astrocyte activation markers in hippocampus from *Aldh1l1-Cre+;Bmal^f/f^* mice vs Cre- controls. All data represent mean+SEM. *p<0.05, **p<0.01, ***p<0.001, ****p<0.0001 by 2-tailed T-tests (A), (E), and (F) or 1-way ANOVA (D) with Hom-Sidak correction for multiple comparisons when applicable.

**Supplemental Figure 3:**
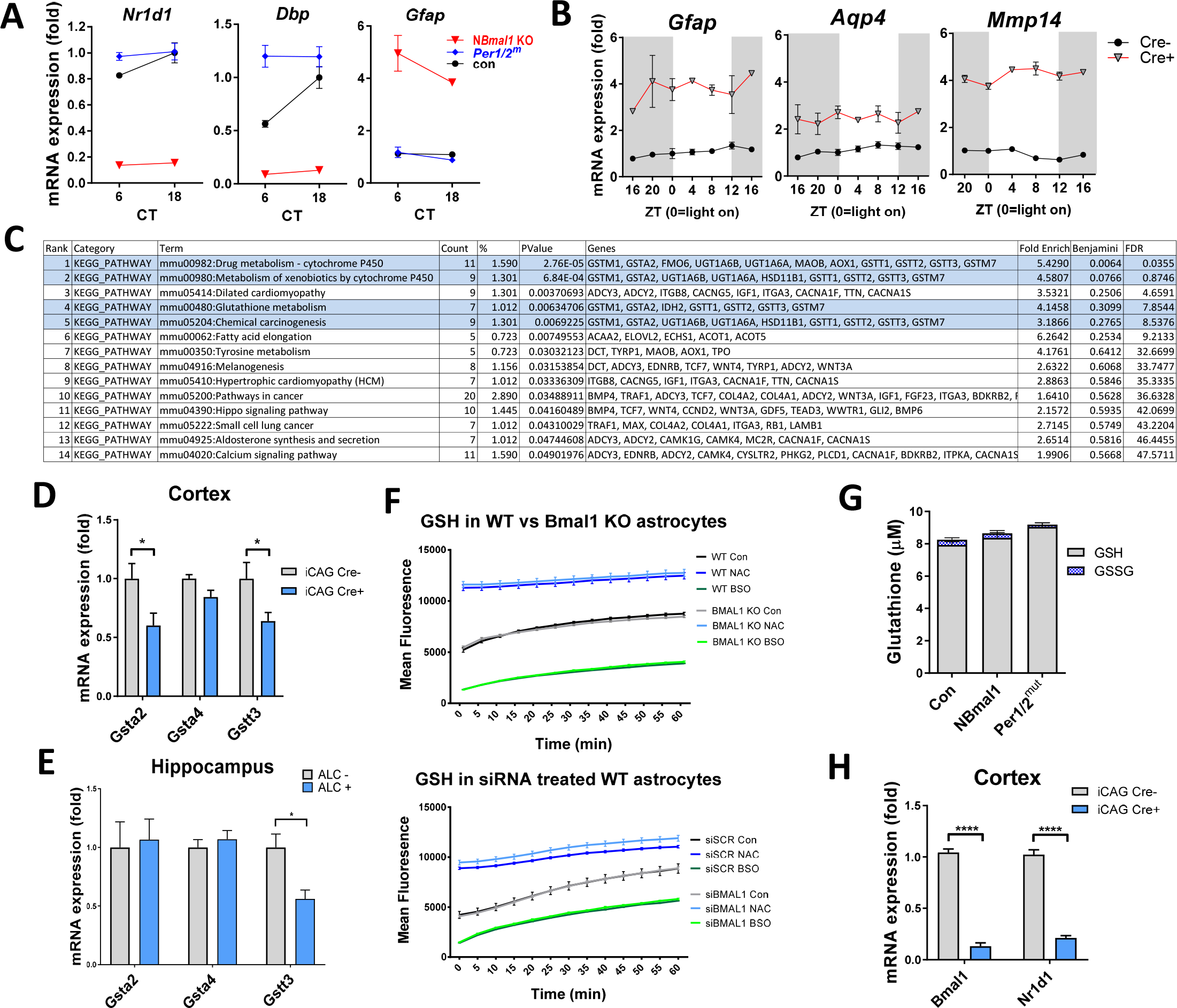
Pathway analysis, GSTx, and GSH with *Bmal1* deficiency. A. Expression of BMAL1 transcriptional targets and *Gfap* from microarray in Fig. 3A at CT 6 and 18. B. Expression of astrocyte activation transcripts *Gfap*, *Aqp4*, and *Mmp14* in cortex of NBmal1 KO (Cre+, red) and Cre-control mice (black) at 4 hour timepoints across the circadian cycle. No clear circadian oscillations were observed in Cre- or Cre+ mice. N=3-4 mice/genotype/timepoint. C. KEGG pathway analysis of NBmal1 KO microarray data. Blue = pathways related to glutathione homeostasis. D. qPCR of mRNA expression of glutathione transferases in the cortex of *iCAG-Cre+;Bmal1^f/^*f mice (Fig. 3F), 9 days after tamoxifen treatment, normalized to Cre-, *Bmal^f/f^* controls. E. qPCR of mRNA expression of glutathione transferases in hippocampus of *Aldh1l1-Cre+;Bmal1^f/f^* mice (Fig. 2H). F. GSH levels in Bmal1 KO (top) and siBmal1 treated WT (bottom) astrocyte enriched cultures along with appropriate controls as measured by fluorometric intracellular glutathione detection assay. G. Quantification of GSH and GSSG levels in cortical tissue (CT 10) from control, NBmal1 KO, and *Per1/2^mut^* mice. None of the averages were significantly different. H. mRNA expression of *Bmal1* and *Nr1d1* in the cortex of *iCAG-Cre+;Bmal1^f/^*f mice used in Fig. 3G, 9 days after tamoxifen treatment, normalized to Cre-, *Bmal^f/f^* controls. All data represent mean±SEM. *p<0.05, ****p<0.001 by 2-tailed T-test with Holm-Sidak correction for multiple comparisons when applicable.

**Supplemental Figure 4.**
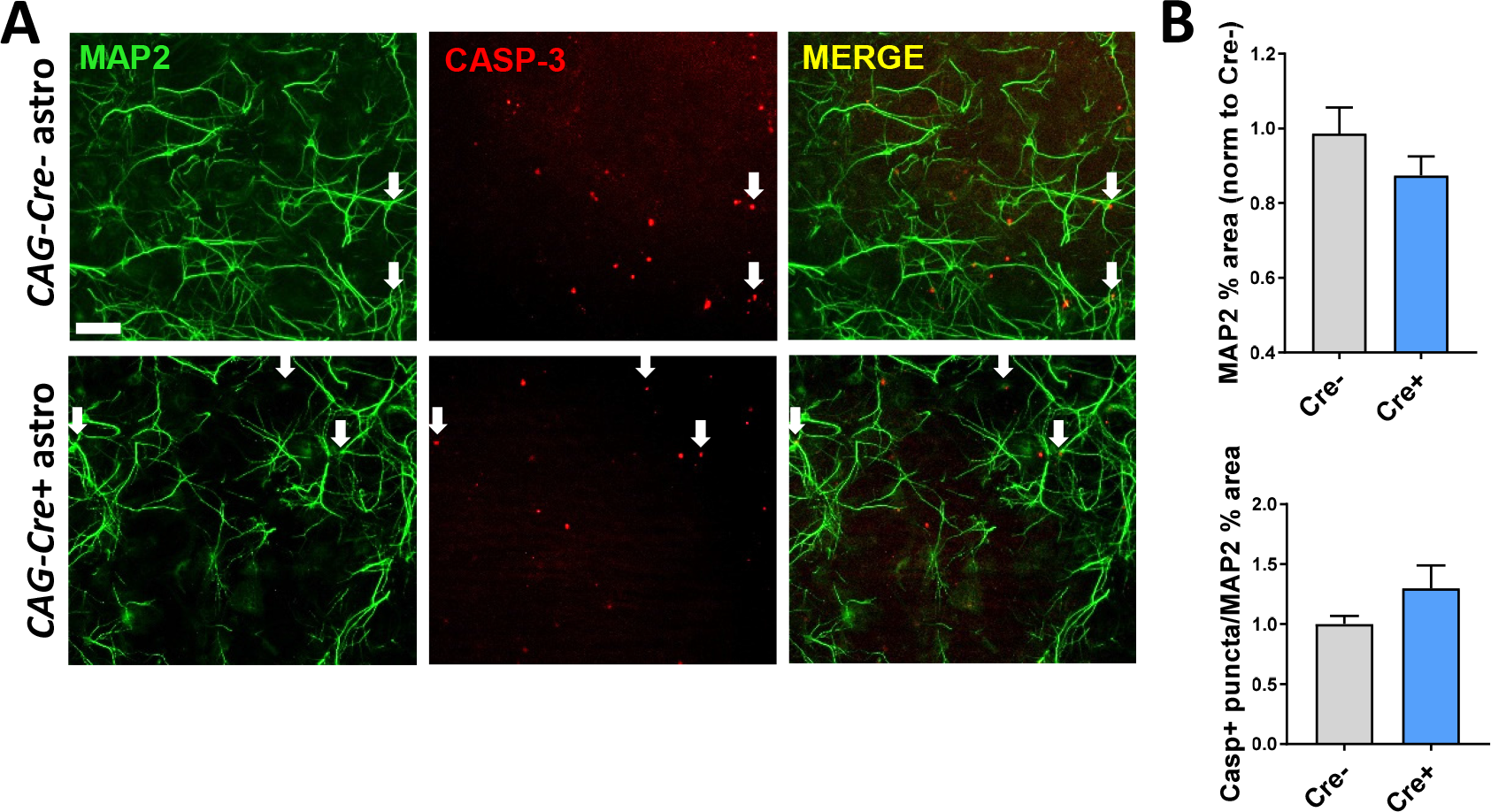
Further characterization of neuronal support and plaque clustering deficits observed with loss of *Bmal1*. A. Representative images showing MAP2+ (green) and cleaved-Caspase-3+ (CASP-3, red, arrows) WT neurons (grown as in Fig. 4A) at DIV 7. Scale bar = 50μm. B. Quantification of MAP2 percent area (top) and cleaved-Caspase-3+ (bottom) neurons from (C) shows non-significant trends toward decreased MAP2 and increased cleaved-Caspase-3+ neurons when neurons are plated on BMAL1-deficient astrocytes (Cre+). All data represent mean+SEM.

## REFERENCES

Barca-Mayo, O., Pons-Espinal, M., Follert, P., Armirotti, A., Berdondini, L., and De Pietri Tonelli, D. (2017). Astrocyte deletion of Bmal1 alters daily locomotor activity and cognitive functions via GABA signalling. Nat Commun 8, 14336.

Bass, J., and Takahashi, J.S. (2010). Circadian integration of metabolism and energetics. Science 330, 1349–1354.

Beaule, C., Swanstrom, A., Leone, M.J., and Herzog, E.D. (2009). Circadian modulation of gene expression, but not glutamate uptake, in mouse and rat cortical astrocytes. PLoS One 4, e7476.

Boisvert, M.M., Erikson, G.A., Shokhirev, M.N., and Allen, N.J. (2018). The Aging Astrocyte Transcriptome from Multiple Regions of the Mouse Brain. Cell Rep 22, 269–285.

Bortell, N., Basova, L., Semenova, S., Fox, H.S., Ravasi, T., and Marcondes, M.C. (2017). Astrocyte-specific overexpressed gene signatures in response to methamphetamine exposure in vitro. J Neuroinflammation 14, 49.

Brancaccio, M., Patton, A.P., Chesham, J.E., Maywood, E.S., and Hastings, M.H. (2017). Astrocytes Control Circadian Timekeeping in the Suprachiasmatic Nucleus via Glutamatergic Signaling. Neuron 93, 1420–1435 e1425.

Breen, D.P., Vuono, R., Nawarathna, U., Fisher, K., Shneerson, J.M., Reddy, A.B., and Barker, R.A. (2014). Sleep and circadian rhythm regulation in early Parkinson disease. JAMA Neurol 71, 589–595.

Chung, W.S., Clarke, L.E., Wang, G.X., Stafford, B.K., Sher, A., Chakraborty, C., Joung, J., Foo, L.C., Thompson, A., Chen, C., et al. (2013). Astrocytes mediate synapse elimination through MEGF10 and MERTK pathways. Nature 504, 394–400.

Clarke, L.E., Liddelow, S.A., Chakraborty, C., Munch, A.E., Heiman, M., and Barres, B.A. (2018). Normal aging induces A1-like astrocyte reactivity. Proc Natl Acad Sci U S A 115, E1896–E1905.

Cronin, P., McCarthy, M.J., Lim, A.S.P., Salmon, D.P., Galasko, D., Masliah, E., De Jager, P.L., Bennett, D.A., and Desplats, P. (2017). Circadian alterations during early stages of Alzheimer’s disease are associated with aberrant cycles of DNA methylation in BMAL1. Alzheimers Dement 13, 689–700.

Curtis, A.M., Fagundes, C.T., Yang, G., Palsson-McDermott, E.M., Wochal, P., McGettrick, A.F., Foley, N.H., Early, J.O., Chen, L., Zhang, H., et al. (2015). Circadian control of innate immunity in macrophages by miR-155 targeting Bmal1. Proc Natl Acad Sci U S A 112, 7231–7236.

Dmytriyeva, O., Pankratova, S., Owczarek, S., Sonn, K., Soroka, V., Ridley, C.M., Marsolais, A., Lopez-Hoyos, M., Ambartsumian, N., Lukanidin, E., et al. (2012). The metastasis-promoting S100A4 protein confers neuroprotection in brain injury. Nat Commun 3, 1197.

Gomez-Nicola, D., Pallas-Bazarra, N., Valle-Argos, B., and Nieto-Sampedro, M. (2010). CCR7 is expressed in astrocytes and upregulated after an inflammatory injury. J Neuroimmunol 227, 87–92.

Grek, C.L., Zhang, J., Manevich, Y., Townsend, D.M., and Tew, K.D. (2013). Causes and consequences of cysteine S-glutathionylation. J Biol Chem 288, 26497–26504.

Hatfield, C.F., Herbert, J., van Someren, E.J., Hodges, J.R., and Hastings, M.H. (2004). Disrupted daily activity/rest cycles in relation to daily cortisol rhythms of home-dwelling patients with early Alzheimer’s dementia. Brain 127, 1061–1074.

Hoyaux, D., Boom, A., Van den Bosch, L., Belot, N., Martin, J.J., Heizmann, C.W., Kiss, R., and Pochet,R. (2002). S100A6 overexpression within astrocytes associated with impaired axons from both ALS mouse model and human patients. J Neuropathol Exp Neurol 61, 736–744.

Iram, T., Ramirez-Ortiz, Z., Byrne, M.H., Coleman, U.A., Kingery, N.D., Means, T.K., Frenkel, D., and El Khoury, J. (2016). Megf10 Is a Receptor for C1Q That Mediates Clearance of Apoptotic Cells by Astrocytes. J Neurosci 36, 5185–5192.

Izumo, M., Pejchal, M., Schook, A.C., Lange, R.P., Walisser, J.A., Sato, T.R., Wang, X., Bradfield, C.A., and Takahashi, J.S. (2014). Differential effects of light and feeding on circadian organization of peripheral clocks in a forebrain Bmal1 mutant. Elife 3.

Kang, S.S., Ren, Y., Liu, C.C., Kurti, A., Baker, K.E., Bu, G., Asmann, Y., and Fryer, J.D. (2017). Lipocalin-2 protects the brain during inflammatory conditions. Mol Psychiatry.

Koike, N., Yoo, S.H., Huang, H.C., Kumar, V., Lee, C., Kim, T.K., and Takahashi, J.S. (2012). Transcriptional architecture and chromatin landscape of the core circadian clock in mammals. Science 338, 349–354.

Kraft, A.D., Johnson, D.A., and Johnson, J.A. (2004). Nuclear factor E2-related factor 2-dependent antioxidant response element activation by tert-butylhydroquinone and sulforaphane occurring preferentially in astrocytes conditions neurons against oxidative insult. J Neurosci 24, 1101–1112.

Li, W., Poteet, E., Xie, L., Liu, R., Wen, Y., and Yang, S.H. (2011). Regulation of matrix metalloproteinase 2 by oligomeric amyloid beta protein. Brain Res 1387, 141–148.

Lian, H., Yang, L., Cole, A., Sun, L., Chiang, A.C., Fowler, S.W., Shim, D.J., Rodriguez-Rivera, J., Taglialatela, G., Jankowsky, J.L., et al. (2015). NFkappaB-activated astroglial release of complement C3 compromises neuronal morphology and function associated with Alzheimer’s disease. Neuron 85, 101–115.

Liddelow, S.A., Guttenplan, K.A., Clarke, L.E., Bennett, F.C., Bohlen, C.J., Schirmer, L., Bennett, M.L., Munch, A.E., Chung, W.S., Peterson, T.C., et al. (2017). Neurotoxic reactive astrocytes are induced by activated microglia. Nature 541, 481–487.

Macauley, S.L., Pekny, M., and Sands, M.S. (2011). The role of attenuated astrocyte activation in infantile neuronal ceroid lipofuscinosis. J Neurosci 31, 15575–15585.

Mailloux, R.J., and Treberg, J.R. (2016). Protein S-glutathionlyation links energy metabolism to redox signaling in mitochondria. Redox Biol 8, 110–118.

Marpegan, L., Swanstrom, A.E., Chung, K., Simon, T., Haydon, P.G., Khan, S.K., Liu, A.C., Herzog, E.D., and Beaule, C. (2011). Circadian regulation of ATP release in astrocytes. J Neurosci 31, 8342–8350.

Martinez-Garcia, A., Sastre, I., Recuero, M., Aldudo, J., Vilella, E., Mateo, I., Sanchez-Juan, P., Vargas,T., Carro, E., Bermejo-Pareja, F., et al. (2010). PLA2G3, a gene involved in oxidative stress induced death, is associated with Alzheimer’s disease. J Alzheimers Dis 22, 1181–1187.

Mohawk, J.A., Green, C.B., and Takahashi, J.S. (2012). Central and Peripheral Circadian Clocks in Mammals. Annu Rev Neurosci 35, 445–462.

Morton, A.J., Wood, N.I., Hastings, M.H., Hurelbrink, C., Barker, R.A., and Maywood, E.S. (2005). Disintegration of the sleep-wake cycle and circadian timing in Huntington’s disease. J Neurosci 25, 157–163.

Musiek, E.S., Bhimasani, M., Zangrilli, M.A., Morris, J.C., Holtzman, D.M., and Ju, Y.E. (2018). Circadian Rest-Activity Pattern Changes in Aging and Preclinical Alzheimer Disease. JAMA Neurol.

Musiek, E.S., and Holtzman, D.M. (2016). Mechanisms linking circadian clocks, sleep, and neurodegeneration. Science 354, 1004–1008.

Musiek, E.S., Lim, M.M., Yang, G., Bauer, A.Q., Qi, L., Lee, Y., Roh, J.H., Ortiz-Gonzalez, X., Dearborn, J.T., Culver, J.P., et al. (2013). Circadian clock proteins regulate neuronal redox homeostasis and neurodegeneration. J Clin Invest 123, 5389–5400.

Nakazato, R., Kawabe, K., Yamada, D., Ikeno, S., Mieda, M., Shimba, S., Hinoi, E., Yoneda, Y., and Takarada, T. (2017). Disruption of Bmal1 Impairs Blood-Brain Barrier Integrity via Pericyte Dysfunction. J Neurosci 37, 10052–10062.

Naoi, M., Maruyama, W., Yi, H., Yamaoka, Y., Shamoto-Nagai, M., Akao, Y., Gerlach, M., Tanaka, M., and Riederer, P. (2008). Neuromelanin selectively induces apoptosis in dopaminergic SH-SY5Y cells by deglutathionylation in mitochondria: involvement of the protein and melanin component. J Neurochem 105, 2489–2500.

Pekny, M., Pekna, M., Messing, A., Steinhauser, C., Lee, J.M., Parpura, V., Hol, E.M., Sofroniew, M.V., and Verkhratsky, A. (2016). Astrocytes: a central element in neurological diseases. Acta Neuropathol 131, 323–345.

Prolo, L.M., Takahashi, J.S., and Herzog, E.D. (2005). Circadian rhythm generation and entrainment in astrocytes. J Neurosci 25, 404–408.

Rathke-Hartlieb, S., Budde, P., Ewert, S., Schlomann, U., Staege, M.S., Jockusch, H., Bartsch, J.W., and Frey, J. (2000). Elevated expression of membrane type 1 metalloproteinase (MT1-MMP) in reactive astrocytes following neurodegeneration in mouse central nervous system. FEBS Lett 481, 227–234.

Schildge, S., Bohrer, C., Beck, K., and Schachtrup, C. (2013). Isolation and culture of mouse cortical astrocytes. J Vis Exp.

Sofroniew, M.V. (2014). Astrogliosis. Cold Spring Harb Perspect Biol 7, a020420.

Song, H., Moon, M., Choe, H.K., Han, D.H., Jang, C., Kim, A., Cho, S., Kim, K., and Mook-Jung, I. (2015). Abeta-induced degradation of BMAL1 and CBP leads to circadian rhythm disruption in Alzheimer’s disease. Mol Neurodegener 10, 13.

Srinivasan, R., Lu, T.Y., Chai, H., Xu, J., Huang, B.S., Golshani, P., Coppola, G., and Khakh, B.S. (2016). New Transgenic Mouse Lines for Selectively Targeting Astrocytes and Studying Calcium Signals in Astrocyte Processes In Situ and In Vivo. Neuron 92, 1181–1195.

Stevanovic, K., Yunus, A., Joly-Amado, A., Gordon, M., Morgan, D., Gulick, D., and Gamsby, J. (2017). Disruption of normal circadian clock function in a mouse model of tauopathy. Exp Neurol 294, 58–67.

Tranah, G.J., Blackwell, T., Stone, K.L., Ancoli-Israel, S., Paudel, M.L., Ensrud, K.E., Cauley, J.A., Redline, S., Hillier, T.A., Cummings, S.R., and Yaffe, K. (2011). Circadian activity rhythms and risk of incident dementia and mild cognitive impairment in older women. Ann Neurol 70, 722–732.

Tso,C.F., Simon, T., Greenlaw, A.C., Puri, T., Mieda, M., and Herzog, E.D. (2017). Astrocytes Regulate Daily Rhythms in the Suprachiasmatic Nucleus and Behavior. Curr Biol 27, 1055–1061.

Wang, T.A., Yu, Y.V., Govindaiah, G., Ye, X., Artinian, L., Coleman, T.P., Sweedler, J.V., Cox, C.L., and Gillette, M.U. (2012). Circadian rhythm of redox state regulates excitability in suprachiasmatic nucleus neurons. Science 337, 839–842.

Wicher, G., Wallenquist, U., Lei, Y., Enoksson, M., Li, X., Fuchs, B., Abu Hamdeh, S., Marklund, N., Hillered, L., Nilsson, G., and Forsberg-Nilsson, K. (2017). Interleukin-33 Promotes Recruitment of Microglia/Macrophages in Response to Traumatic Brain Injury. J Neurotrauma.

Wyse, C.A., and Coogan, A.N. (2010). Impact of aging on diurnal expression patterns of CLOCK and BMAL1 in the mouse brain. Brain Res 1337, 21–31.

Xiao, Q., Yan, P., Ma, X., Liu, H., Perez, R., Zhu, A., Gonzales, E., Burchett, J.M., Schuler, D.R., Cirrito, J.R., et al. (2014). Enhancing astrocytic lysosome biogenesis facilitates Abeta clearance and attenuates amyloid plaque pathogenesis. J Neurosci 34, 9607–9620.

Yamanaka, K., Chun, S.J., Boillee, S., Fujimori-Tonou, N., Yamashita, H., Gutmann, D.H., Takahashi, R., Misawa, H., and Cleveland, D.W. (2008). Astrocytes as determinants of disease progression in inherited amyotrophic lateral sclerosis. Nat Neurosci 11, 251–253.

Yang, G., Chen, L., Grant, G.R., Paschos, G., Song, W.L., Musiek, E.S., Lee, V., McLoughlin, S.C., Grosser, T., Cotsarelis, G., and FitzGerald, G.A. (2016). Timing of expression of the core clock gene Bmal1 influences its effects on aging and survival. Sci Transl Med 8, 324ra316.

Zamanian, J.L., Xu, L., Foo, L.C., Nouri, N., Zhou, L., Giffard, R.G., and Barres, B.A. (2012). Genomic analysis of reactive astrogliosis. J Neurosci 32, 6391–6410.

Zhang, R., Lahens, N.F., Ballance, H.I., Hughes, M.E., and Hogenesch, J.B. (2014a). A circadian gene expression atlas in mammals: Implications for biology and medicine. Proc Natl Acad Sci U S A 111, 16219–16224.

Zhang, Y., Chen, K., Sloan, S.A., Bennett, M.L., Scholze, A.R., O’Keeffe, S., Phatnani, H.P., Guarnieri,P., Caneda, C., Ruderisch, N., et al. (2014b). An RNA-sequencing transcriptome and splicing database of glia, neurons, and vascular cells of the cerebral cortex. J Neurosci 34, 11929–11947.

